# Surface Acoustic Wave Integrated Microfluidics for Repetitive and Reversible Temporary Immobilization of *C. elegans*

**DOI:** 10.1101/2022.06.20.496864

**Authors:** Nakul Sridhar, Apresio Kefin Fajrial, Rachel Doser, Frederic Hoerndli, Xiaoyun Ding

**Affiliations:** Paul M. Rady Department of Mechanical Engineering, University of Colorado, Boulder, Colorado, USA; Department of Biomedical Sciences, Colorado State University, Fort Collins, Colorado, USA; Biomedical Engineering Program, University of Colorado, Boulder, Colorado, USA

## Abstract

*Caenorhabditis elegans* is an important genetic model for neuroscience studies due to its unique combination of genetics, transparency, complete synaptic connectome, and well-characterized behaviors. These factors, in turn, enable analyses of how genes control connectivity, neuronal function, and behavior. To date, however, most studies of neuronal function in *C. elegans* are incapable of performing microscopy imaging with subcellular resolution and behavior analysis in the same set of animals. This constraint stems from the immobilization requirement for high-resolution imaging that is incompatible with behavioral analysis. In particular, conventional immobilization methods often lead to either irreversible, partial, or slowly reversible immobilization of animals preventing a multiplexed approach. Here, we present a novel microfluidic device that uses surface acoustic waves (SAW) as a non-contact method to temporarily immobilize worms for a short period (40 seconds). This device allows non-invasive analysis of swimming behavior and high-resolution synaptic imaging in the same animal. In addition, because of the low impact of this SAW approach, the device enables fast, repeated imaging of single neurons and behavior in the same animals for three to four days. We anticipate that this device will enable longitudinal analysis of animal motility and subcellular morphological changes during development and ageing in *C. elegans*.

## Introduction

The nematode *Caenorhabditis elegans* serves as a powerful tool to study a large range of biological processes relevant to human health, including responses to external stimuli, cellular and subcellular development, and the pathology of numerous diseases^1–7^. Many of these studies require immobilization of the *C. elegans* for high resolution imaging and analysis. The most popular methods for immobilizing worms include administering chemical anesthetics^8–9^ or using polystyrene nanoparticles on agarose pads slowing animals by friction^11^. These methods are effective at preventing worms from moving, but often lead to slow or no recovery of worm behavior and can lead to either physical damage or unwanted secondary signaling effects. Therefore, longitudinal studies of behavior and cellular microscopy in the same animal have not been feasible. Common developmental and ageing studies applied correlative association of imaging and behavior or circuit activity analysis, involving different set of animals for each respective assay^12–14^. Despite groups of *C. elegans* being isogenic, the ability to perform behavior and microscopy within the same set of animals will improve our ability to correlate cellular function and behavior, distinguishing between individual variation in response to environmental manipulation and experimental variations. Currently, an immobilization technique that can facilitate this multiplex study design is lacking.

The size of *C. Elegans* and their ability to live for several days in liquid environments^15^ have made them ideally suited for manipulation by lab-on-a-chip microfluidic devices. The transparent gas permeable and precise micrometer molding of the silicone PDMS allow the engineering of microfluidic devices that control the placement, environment, and behavior of *C. elegans* animals^16–19^. Several devices have been previously described allowing for the controlled, reversible immobilization of *C. elegans* animals. Some are mechanical in nature such as the one described by Hulme *et al.,* which used an array of tapered channels that used the pressure difference between the inlet and outlet to physically immobilize worms in the chamber, which could then be released by reversing the pressure^20^. Another mechanical method consists of pressurizing a flexible PDMS membrane to compress the microfluidic channel, therefore trapping animals temporarily^21–23^. Mechanical immobilization typically requires specifics size constraints and must be adjusted depending on the age or average size of C. elegans animals used. Alternatively, Chokski *et al* introduced CO_2_ into a worm chamber that could reversibly immobilize single worms up to one hour^24^. However, hypoxia^25^ has been shown to induce molecular stress signaling in neurons which could be a problem in studies of neurodegeneration and ageing. Heating or cooling the animals’ environment has also been shown to temporarily immobilize animals. Heating a microfluidic chamber to a specific knockdown temperature range has been shown to introduce quiescence in animals^26^. This approach required a complex integration of laser and electrical systems. Similarly, cooling a chamber could also reduce worm movement enough to allow high resolution imaging^27–28^. Yet, this cooling process to reach sufficient immobilization temperature can be time consuming. Therefore, a simple non-contact method for fast reversible immobilization of animals is necessary to harness the potential of *C. elegans* in developmental and ageing studies.

Surface acoustic wave (SAW) technology integrated with microfluidic devices have gained more traction for a variety of biological applications^29–30^. SAW microfluidic devices have previously been used to apply forces to *C. elegans* to precisely move^31^, rotate^32^, and perturb their chemosensory learning^33^. Here, we introduce a novel microfluidic device that can apply SAW to reversibly immobilize a single *C. elegans* for up to 30 seconds, allowing for high resolution images up to the synaptic level. Furthermore, the device enabled several rounds of immobilization followed by free swimming over a period of 3-4 days, enabling longitudinal imaging of individual synapses and swimming behavior in the same animal. The immobilization process is gentle and non-invasive, enabling rapid behavioral recovery immediately after the acoustic wave is turned off. Altogether, this technology will pave the way toward longitudinal and multiplexed analyses of C. elegans.

## Experimental Methods

### Device Design

The schematic of our SAW worm immobilization device is shown in **Fig. 1A**. The device consists of rectangular polydimethylsiloxane (PDMS) chamber filled with fluid bonded to a lithium niobate substrate (**Fig. 1B**). The chamber has a height of 60 μm and a width of 500 μm. 200 μm channels connect the chamber to the inlet and outlet. An interdigital transducer (IDT) is fabricated atop the substrate adjacent to the PDMS chamber. When a radio frequency (RF) signal (50 MHz) is input into the IDT, a traveling wave is generated and propagates along the top of the substrate towards the PDMS chamber (**Fig. 1C**). As the waves reach the fluid in the chamber, the difference in velocity of acoustic propagation along the fluid versus the substrate causes some of the wave energy to leak into the fluid. This leakage propagates as an acoustic radiation force that can be precisely controlled via the power of the RF signal input^28^.

**Figure 1:**
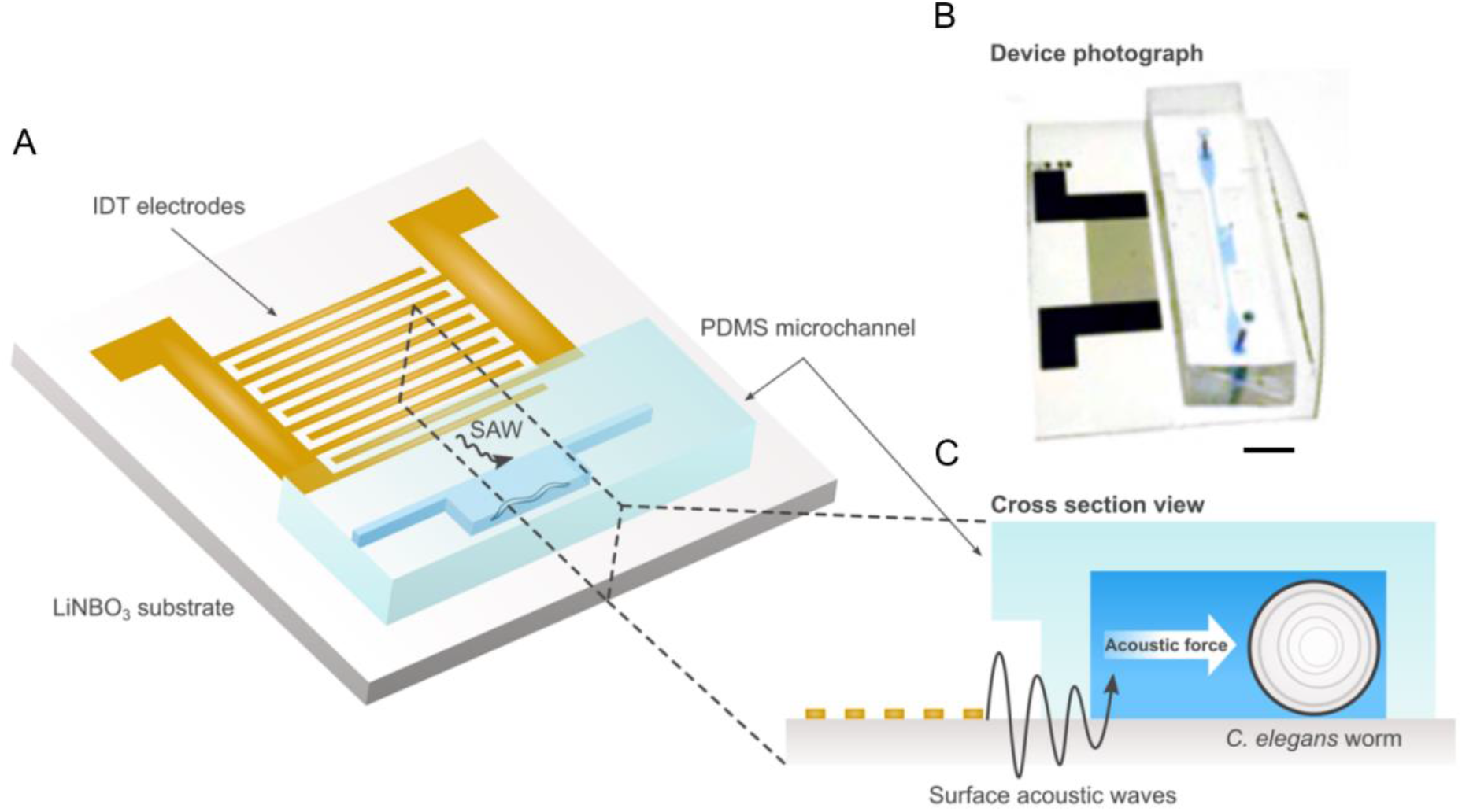
**A,** Schematic and **B,** photo of SAW device illustrating single IDT adjacent to PDMS worm chamber. Scale bar: 5 mm. **C,** Cross-section view of device showing IDT producing traveling SAW acoustic force into chamber to immobilize worms.

### Device Fabrication

The immobilization device consists of the lithium niobate substrate patterned with a single IDT, and the PDMS chamber. The LiNbO_3_ substate was patterned onto a 3inch, Y+36°X-propagation LiNbO_3_ wafer using standard photolithography techniques. First, a layer of photoresist (S1813, Dow, USA) was spin coated onto the wafer. The IDT structures were then patterned onto the wafer using UV light exposure and then developed using a photoresist developer (MF319, Dow, USA). Next, layers of chrome and gold (Cr/Au, 10/100nm) were deposited using e-beam evaporation. Excess photoresist was removed using a lift-off process (Remover PG, Kayaku, Japan). Finally, the wafer was diced into individual devices each containing a single IDT. Each individual IDT consists of 20 electrode pairs with a spacing of 20 μm and an aperture of 6 mm.

The PDMS chamber was made using a negative replica from an SU8 master mold. To fabricate the SU8 mold, a 60 μm thick layer of SU8 2025 photoresist (MicroChem, USA) was spin-coated on a 3-inch silicon wafer. The wafer was then patterned using standard optical lithography. To create PDMS replicas, the master mold surface was first modified by placing it in a silane vapor for 30 minutes. Then, a mixture of PDMS base and cross-linker (Sylgard 184, Dow Corning, USA) with a ratio of 10:1 w/w was poured onto the master mold and cured at 65°C for one hour. Inlet and outlet holes with a 0.75 mm diameter, as well as a 0.35 mm diameter hole in the center of the chamber to measure the temperature in control devices, were punched using a biopsy hole puncher. Finally, the PDMS chamber was bonded to the LiNbO3 substrate using oxygen plasma (PDC-001, Harrick Plasma, USA). To ensure a strong bond, the device was baked overnight at 65°C.

### Strains and maintenance

*C. elegans* strains were cultured and maintained at 20°C under standard conditions^34^. The initial characterization of the immobilization device and the longitudinal assays was completed using wild-type N2 Bristol strains. Animals used for fluorescent imaging of neurons and synapses imaging either contained the integrated multicopy array *nuIs25* (KP1148) or *akIs141* (FJH18). *nuIs25* and *akIs141* both express the *C. elegans* ionotropic glutamate receptor homologue GLR-1 tagged with GFP either under its native *GLR-1p^35^* or AVA specific *Rig-3p* promoters^36^. Initially, animals used for fluorescence imaging contained the *lite-1(ok530)* mutation, RB765 strain to prevent light-induced side effects. The ST66 or *(ncIs17 [hsp16.2p::eGFP + pBluescript])* strain was used to measure the biological effect of SAW induced-heat (supplemental figure 3).

### Device Operation

An RF function generator (N5171B, Keysight, USA) and power amplifier (403LA, E&I, USA) were used to input the signal into the IDTs. To identify the resonant frequency of the devices, we used a network analyzer (E5061B, Keysight, USA) in combination with visually tracking the velocity of polystyrene microparticles in the fluid that were pushed by the acoustic radiation force. Depending on the individual device, the resonant frequency was identified as approximately 46 MHz.

Initial characterization of the immobilization device was conducted on an inverted microscope (Eclipse Ti2, Nikon, Japan). Worms in the chamber were imaged with a digital CMOS camera (Orca-Flash 4.0, Hamamatsu, Japan) in combination with the included imaging software (HCImage Live, Hamamatsu, Japan). GFP-B fluorescent filter cubes (495 nm, ET 49002, Chroma, USA) were used to image fluorescent neurons in high resolution. Tracking the worm movement and further processing of images and videos were completed using ImageJ^37^ For measuring the temperature inside the chamber, a digital thermocouple was used to track the change when SAW was applied.

For all worm experiments, synchronized animals were allowed to grow until the Adult day 1 stage in normal NGM plates. They were then washed off the NGM plates with M9 buffer, and a single worm was manually transferred into the immobilization device using a 10 μL micropipette.

### Immobilization characterization

The locomotion of *C. elegans* during SAW application was quantified by visually tracking the movement at the tip of the head of the animal inside the device. The Manual Tracking plugin^38^ in ImageJ was used to follow the tip of the head and calculate the distance moved at each frame. A worm was defined as immobilized when no noticeable head movement could be tracked over >5 frames.

For each test, 3 cycles of SAW were applied so that the animal was immobilized for 30 seconds. Between each cycle, there was a 3-minute recovery period where the worm was allowed to swim freely in the chamber. The head movement of the worm was tracked as the SAW was applied, as well as after the SAW was turned off to quantify recovery. Various SAW powers and duty cycles were tested. A worm was scored as recovered if it exhibited similar swimming behavior 3 minutes after SAW application as seen before SAW application.

### Quantification of longitudinal worm swimming

Experiments were conducted over a 72-hour period. SAW (2W-50% duty cycle immobilization, 1W-50% duty cycle hold) was applied at hour 12, 36 and 60 to temporarily immobilize the worm for 30 seconds. SAW treatments were compared to control animals that were allowed to swim freely inside the chamber. Animal swimming was visually analyzed at hour 0, 24, 48, and 72. Animals were scored as healthy if they exhibited a response to 10 seconds of blue light and were swimming at an average bending frequency greater than 1 Hz. Every 12 hours, the liquid in the chamber, along with any offspring, was replaced with fresh liquid OP50 in an M9 medium. To prevent any evaporation between tests., devices were stored in a high humidity chamber at room temperature. The same individual animals were tracked over multiple days.

### Quantification of longitudinal behavior tracking

Experiments were conducted over a 48-hour period. At hour 0, 24, and 48, worms were tracked for 60 seconds and the average bending frequency over the time frame was calculated. One full bending cycle was defined as a full wave of dorsal-ventral bends through its body^39^. SAW (2W-50% duty cycle immobilization, 1W-50% duty cycle hold) was applied at hour 12 and 36 to temporarily immobilize the worm for 30 seconds. SAW animals were compared against control animals that were allowed to swim freely in the chamber for the same time frame. Feeding and storage methods were the same as previously described for the longitudinal worm swimming assay.

### Longitudinal fluorescent imaging of single neurons

Experiments were conducted over a 48-hour period. At hour 0, 24, and 48, SAW (2W-50% duty cycle immobilization, 1W-50% duty cycle hold) was applied to temporarily immobilize and hold the worm. Once the worm was visually deemed to be immobilized, imaging was switched from brightfield to wide field GFP fluorescence imaging. Single 500 ms exposure wide field images of glutamatergic neurons (including AVA) were taken using an inverted microscope (Eclipse Ti2, Nikon, Japan). Control images were taken of animals treated with sodium azide (50 mM). Since sodium azide treatment compromised animals’ health, different control animals were tracked as representative examples for fluorescent intensity for each day. For the SAW-treated animals, feeding and storage methods were the same as previously described for the longitudinal worm swimming assay. The fluorescence in the soma of the AVA cell bodies was quantified in ImageJ using the measure tool and ellipse tool to draw an ROI tightly constrained around the fluorescence of the cell body image at mid-range of the total intensity. The fluorescence within the ROI around the soma was measured first and then the ROI was displaced in a region of the image without punctate fluorescence used as background. For all measurements this background was subtracted from the cell body fluorescence. Quantification of the AVA cell body fluorescence was done on all images except on those where movement of the animal could be detected in the images.

### Temperature characterization

To measure SAW-induced temperature increase, a thermal probe was inserted into the device from above through the 0.35 mm central inlet. An initial standard room temperature of the fluid of approximately 20 °C was established. Then, temperature was tracked for various initial high power SAW applications, as well as switches to low power SAW for holding the immobilization. After the SAW was turned off, the cool-down temperature was tracked along 15-60 second intervals until room temperature was reached. For pure heating-only control tests, devices were placed upon a glass heating plate. The heating plate was turned on and the temperature was increased until the animal was immobilized. The heating plate was then immediately turned off and the fluid in the device was allowed to cool back to room temperature.

For heat stress quantification, SAW (2W-50% duty cycle immobilization, 1W-50% duty cycle hold) was applied to temporarily immobilize *ST66* animals. Positive control animals were placed in a 32 C hot bath for 1 hour. Negative control animals had no heat applied to them. Afterwards, all animals were allowed to swim in chamber for one hour. Animals were then immobilized with sodium azide and single 500 ms exposure images were then taken of the worm bodies. For each image, the average intensity of the image was quantified in ImageJ by using the measure tool and the ellipse tool to draw an ROI around the entire worm body. The background of all images was initially normalized to an equivalent intensity range.

## Results

### SAW induced rapid animal immobilization and instantaneous recovery

Initially, we visually confirmed the SAW device’s ability to immobilize worms by imaging single brightfield frames with a 100 ms exposure time. While a free-swimming worm produces a blurry image (**Fig. 2A**), we could see a clear outline of the animal when applying SAW (**Fig. 2B**). To evaluate the effectiveness and controllability of the device, we then tracked movement at the tip of the head of an individual worm over 3 cycles of immobilization and recovery (**Fig. 2C** and **Video S1**). The immobilization process, initiated with 2W-50% SAW, typically occurred in 60 seconds. Switching the SAW to lower power (1W-50%), the worm can be temporarily held immobilized for more than 30 second which is sufficient timeframes for various high-resolution imaging needs. When the SAW was turned off, in only a few seconds, the worm started thrashing indicating that the worm can rapidly recover after immobilization process.

**Figure 2:**
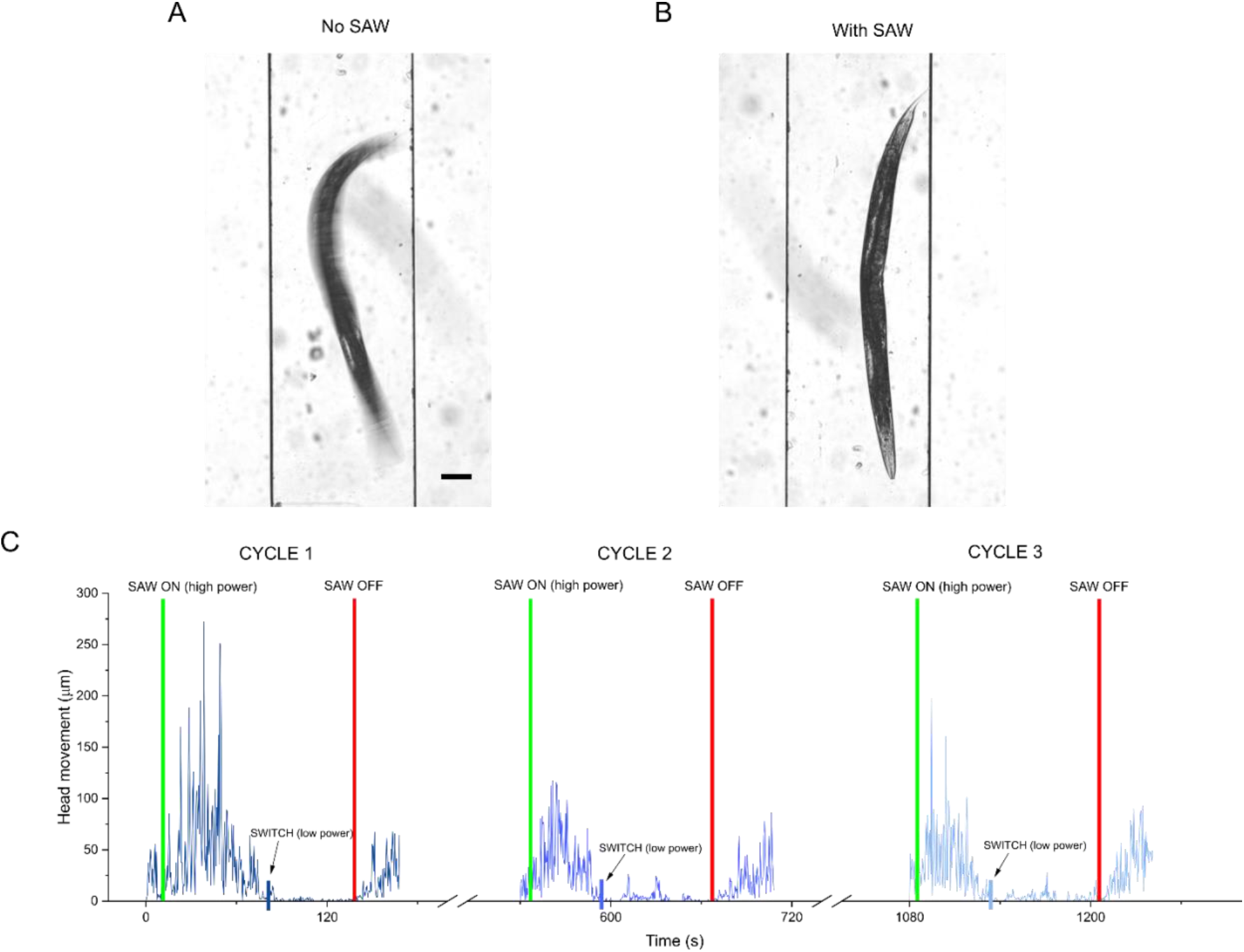
Imaging and characterization of worm immobilization process. Bright field images (100 ms exposure) of single *C. elegans* animal in chamber comparing **A,** free worm swimming when no SAW is applied and **B,** immobilization after SAW is applied. Scale bar: 100 μm. **C,** Head movement of single representative *C. elegans* animal over 3 immobilization and recovery cycles (see methods). Initially, higher power SAW (2W-50% duty cycle) was used to immobilize animals. Once animals stopped moving, SAW was switched to lower power (1W-50%duty cycle) to hold animals in immobilized state for 40 seconds. Green vertical line: high power SAW on. Blue vertical lines: switch SAW to lower power. Red vertical line: SAW off.

We characterized various input powers along with comparing continuous and 50% duty cycles to find the most effective SAW protocol to fully immobilize an animal for a long enough time frame to focus on fluorescent structures of interest and capture images while still allowing worms to recover after the 3 cycles (**Table 1** and **Fig. S1**). We found that the best way to extend the immobilization time without negatively affecting worm health was to apply SAW at two successive power levels. First, we applied higher power SAW to initially immobilize the animals. We then immediately switched to a 50% lower power SAW as a hold step to maintain the immobilized state. Keeping the high-power SAW on to hold immobilization might result in decreasing worms’ viability. Meanwhile, simply turning SAW off without the hold step could only keep animals immobilized for approximately 10 seconds. We compared different combinations of high-power SAW (2 W continuous, 1 W continuous, 2 W-50% duty cycle) and low power SAW (0.5 W continuous, 1W-50% duty cycle). We found that the best result was achieved using an approach combining both high-power and low-power steps. Thus, to achieve a long immobilization for imaging while still allowing healthy recovery, we used a 2 W – 50% duty cycle, followed shortly by a 1 W – 50% duty cycle which we designated as our standard protocol and used for all further experiments. Using this combinatorial approach, we were consistently able to hold worms immobilized for over 30 seconds for 3 cycles with 90% of animals swimming at the same rate as prior to SAW immobilization (**Table 1**). Applying high-power SAW until immobilization (between 30-70 seconds depending on the individual animal), then immediately switching to low-power SAW for 30 seconds using the standard protocol was selected as a safe immobilization length that allowed us enough time to fluorescently image animals under high resolution while still maintaining worm health over multiple cycles.

**Table 1:** Swimming worms after 3 immobilization/recovery cycles for different SAW protocols

| **Power**<br>**(High power/low power,**<br>**duty cycle)** | **Worms**<br>**tested** | **Swimming**<br>**worms after 3**<br>**cycles** | **Percent**<br>**swimming** | **Max**<br>**immobilization**<br>**hold – Cycle 1 (s)** |
| --- | --- | --- | --- | --- |
| **2W/1W, 50% duty cycle** | **41** | **a37\*\*\*** | **90.2** | **b39.13\*\*\*** |
| **2W/-- continuous** | 14 | 10<sup>NS</sup> | 71.4 | 8.68 |
| **1W/-- continuous** | 12 | 10<sup>NS</sup> | 83.3 | 6.84 |
| **2W/0.5W continuous** | 10 | 5 | 50.0 | 14.98<sup>NS</sup> |
| **1W/0.5W continuous** | 10 | 4 | 40.0 | 26.17<sup>NS</sup> |

<sup>a</sup> \*\*\*P< 0.001, chi-square test of independence. NS = no significance.
<sup>b</sup> \*\*\*P< 0.001, two-sided, unpaired t-test. NS = no significance.

### SAW immobilization did not trigged heat stress on animals

Generally, it is accepted that sustained exposure to temperatures greater than 25° C has a negative effect on *C. elegans* health, reproduction, and lifespan^40–42^. However, a previous study by Chuang *et al* showed that short term exposure (seconds to minutes) to elevated temperatures in the so-called heat knockdown range of 31-37 °C will induce a temporary shutdown of neural function in worms^27^. Once the temperature is lowered, regular neural functions resumed. This effect was shown to be reversible for multiple cycles. Furthermore, there was no significant effect on worm lifespan and progeny^27^.

SAW produces a known heating effect along with the acoustic force. We characterized the SAW induced heating of a representative device and showed a temperature increase in the chamber from room temperature (21.7 °C) to 29.1 °C in 30 seconds, 33.1 °C in 60 seconds, and 36.2 °C in 90 seconds when applying high power 2 W-50% duty cycle SAW (**Fig. S2)**. After switching to the lower power hold step (1 W-50% duty cycle) the temperature stabilized at approximately 32 °C (**Fig. S2**). We also measured the minimum temperature at which a worm was immobilized, and found it occurs at 28.1 °C. This is significantly lower than an immobilization temperature of 35 °C recorded when using a heating plate substrate to represent a pure heating effect (**Video S2**). Moreover, this is lower than the previously reported heat knockdown immobilization range of 31-37 °C^27^. We believe the lower immobilization temperature indicates that both acoustic pressure and heat play a key role in the immobilization mechanism of our device, and therefore imparts less environmental stress on animals than just pure heating.

To verify that heat stress was not having a significant effect during the immobilization process, we tested our protocol on the ST66 strain containing *ncIs17[hsp16.2p::eGFP]* expressing a heat-shock inducible GFP marker. We compared up to 3 cycles of SAW immobilization with a positive control, where animals were placed in a 32 °C hot bath for 1 hour, and a negative control, where no heat was applied. One hour after heat or SAW was applied, we captured fluorescent images of animals immobilized with sodium azide (**Fig. S3A).** For the different conditions, we compared the average fluorescent intensity in worm bodies (**Fig. S3B**). The results showed that applying SAW for up to three cycles does not induce any significant heat shock response based on expression of GFP from the *hsp16.2p::eGFP* construct.

### Worm immobilization via SAW facilitated high resolution imaging

We used the developed SAW immobilization protocol to capture high resolution fluorescent images of cell bodies of interest. Using a wide-field microscope, we were able to showcase the dynamic usability of the SAW device, which allowed for rapid switching between an immobilized and free-swimming state. Here, a single *nuIs25* (KP1148) worm was immobilized with SAW and imaged for 3 cycles (**Fig. 3A**). In between each cycle, SAW was turned off and the animal was allowed to swim freely for 3 minutes. We proceeded to image single AVA neurons in *akIs141* (**Fig. 3B**). At 40x magnification, we were able to image single synapses containing GFP tagged GLR-1 receptors at high resolution. (**Fig. 3B**). The images confirmed that our device protocol was able to effectively immobilize animals for sufficient time to focus our camera on the cells of interest and use exposures up to 500 ms, which were required for clarity at higher magnifications.

**Figure 3:**
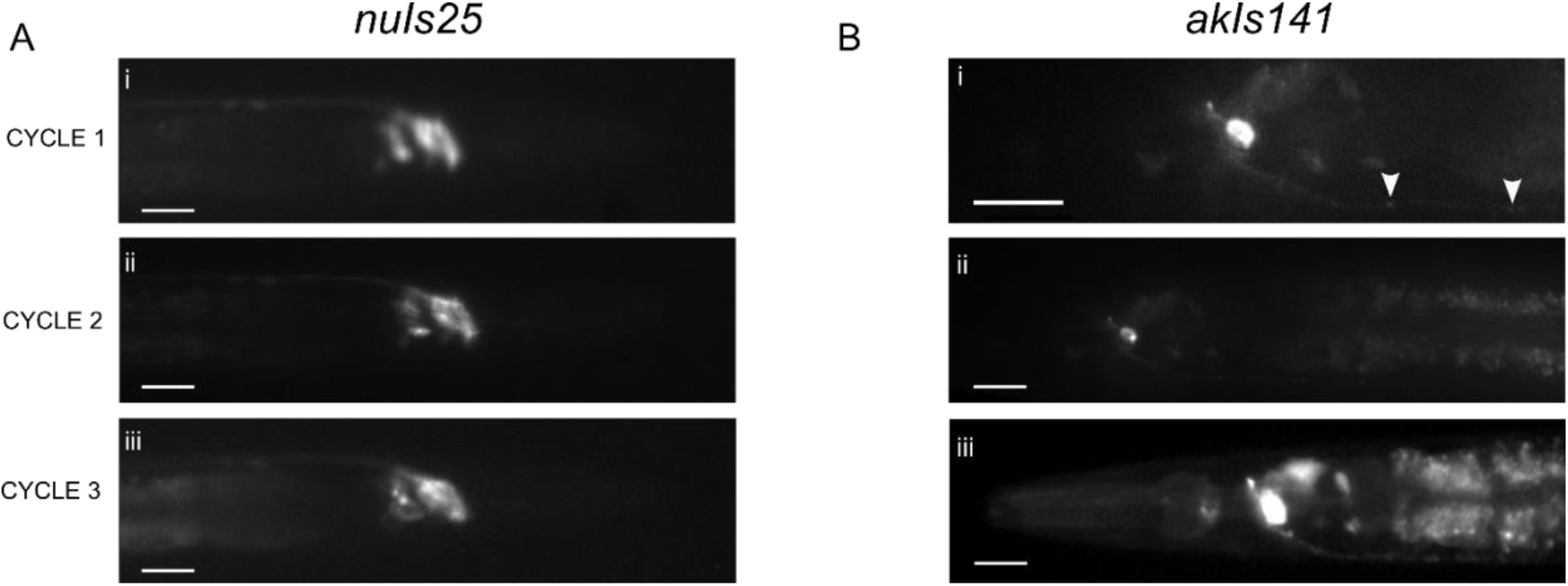
Imaging capability of SAW immobilization device. **A,** Fluorescent images of GFP tagged GLR-1 in (i-iii) a single *nuIs25* animal immobilized with SAW taken over 3 cycles, each 3 minutes apart. The animal was allowed to swim freely in chamber between each cycle. **B,** Wide-field fluorescent images of GFP tagged GLR-1 in a single neuron (AVA) of *akIs141* animals with (i) 40x objective and (ii-iii) 20x objective. Synaptic puncta are visible at 40x (arrowheads). Scale bar: 20μm at 20x, 10μm at 40x.

### SAW immobilization did not affect long term worm survival on-chip

We next tested how animal behavior and neuronal fluorescence would be affected by multiple immobilization recovery cycles during a span of 72 hours in the microfluidics device. The ratio of healthy worms immobilized with SAW once per day was compared against age matched control animals allowed to freely swim in the device for the same time frame (**Fig. 4A**). Worms that responded to 10 seconds of blue light stimulus and swimming at an average bending frequency >1 Hz were deemed as healthy. Over 80% of animals immobilized once a day with SAW survived in the chip over the 72-hour period. The results showed no significant difference in worm health between SAW immobilized and free-swimming control animals, suggesting that SAW application had a negligible impact on long-term worm health. In addition to swimming, a representative animal was imaged under bright field every 24 hours showing no superficial signs of worm stress such as internal organ damage or a clear body (**Fig. 4B**).

**Figure 4:**
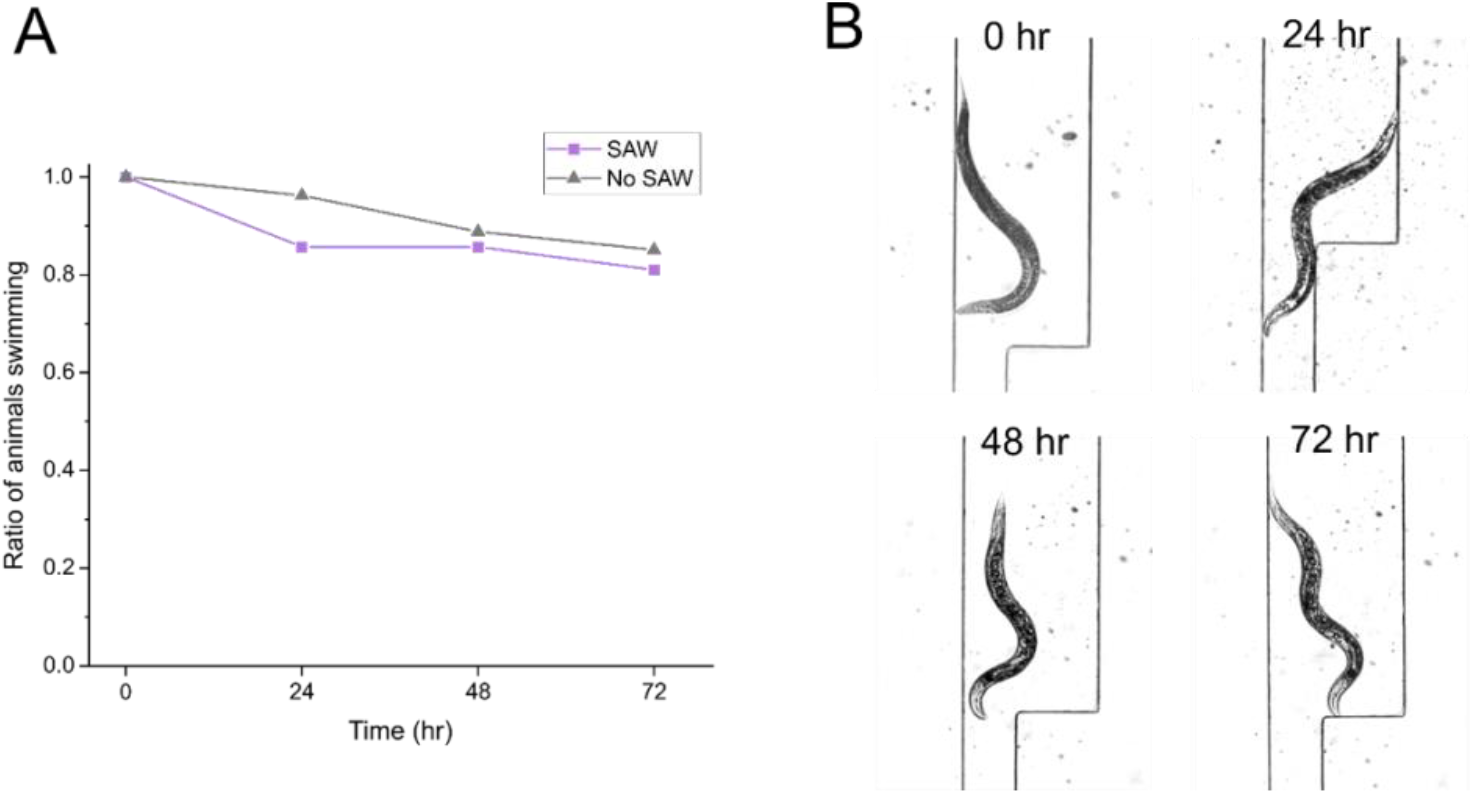
Longitudinal survival on-chip. **A,** Ratio of worm survival in chamber over 72 hours comparing animals that were immobilized with SAW once per day (at hour 12, 36, 48) versus control animals in chamber where no SAW was applied. **B,** Representative bright field image of the same freely swimming animal in chamber imaged at 0, 24, 48, and 72 hours after SAW immobilization once per day. For **A,** at least 20 animals were tracked for each condition.

### Multiplexed tracking of longitudinal swimming behavior and fluorescent imaging achieved using SAW immobilization

Finally, we demonstrated the potential for our SAW device to be used as a platform to track worm behavior and perform fluorescent microscopy imaging in the same animals over multiple days. For the behavior assay, we analyzed animals at 0, 24, and 48 hours and compared the average bending frequency of single worms in the device that were immobilized with SAW and allowed to recover against control animals that could swim freely in the device (**Fig. 5A**). For all time points, there was no significant difference between the bending frequency of the SAW and control worms. Both the SAW and control animals exhibited similar trends in their swimming behavior as they aged. Although there was no significant difference in daily average bending frequency for either the SAW or control populations, individual animals for both the SAW and control tests displayed substantial differences in their swimming behavior each day. This considerable day-to-day change in swimming frequency has been previously reported^43^. On average, SAW worms showed a 38% decrease and control worms showed a 9% decrease at 48 hours. For both SAW and control tests, individual animals showed high variability in movement at each time point. For example, at 24 hours, SAW animals demonstrated bending frequencies between 0.49-1.86 Hz, while control animals frequencies ranged from 0.86-1.47 Hz. This substantial individuality in different animals, despite genetic and environmental homogeneity, has also previously been demonstrated^44^.

**Figure 5:**
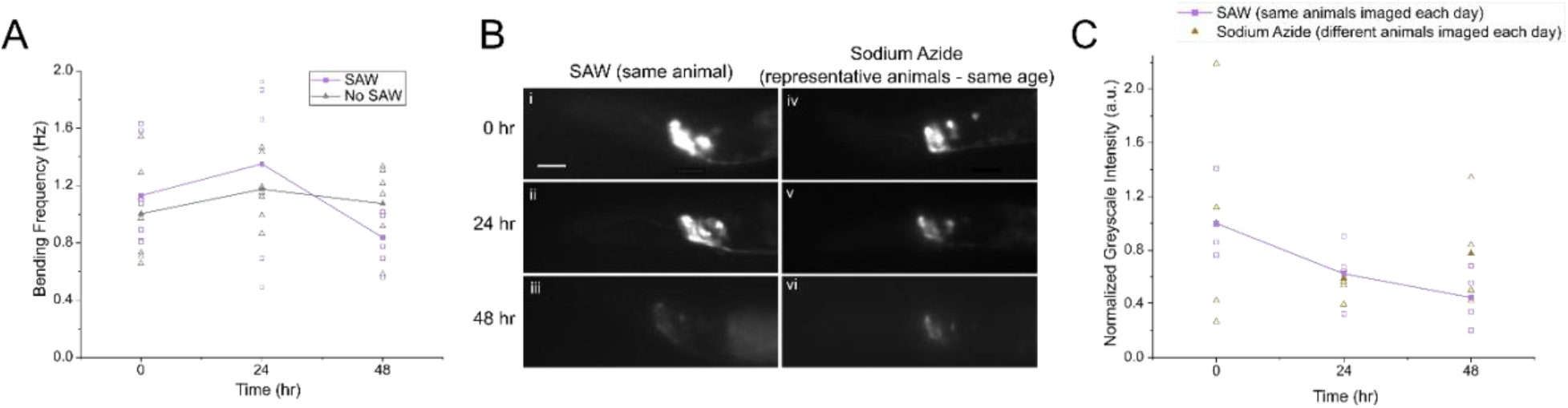
Multiplexed tracking of longitudinal behavior and fluorescence. **A,** Bending frequency of animals at 0, 24, and 48 hours comparing animals immobilized with SAW once per day (at hour 12 and 36) versus control animals freely swimming in chamber. **B,** Representative wide-field fluorescent images of (i-iii) a single *nuIs25* animal immobilized with SAW taken once per day, and (iv-vi) different representative animals immobilized with sodium azide corresponding to the same age. Scale bar: 20μm. **C,** Normalized greyscale intensity in GLR-1::GFP in the soma of the AVA neurons of images taken once a day over 3 days. Note: for animals immobilized with SAW, the same individual animals were imaged over multiple days. For animals immobilized with sodium azide, representative images of different animals for the corresponding age were taken. Data are mean and individual points. For **A,** at least seven animals were tracked at 0, 24, and 48 hours. For **C,** at least four animals were imaged each day for both SAW and sodium azide experiments.

For the fluorescent imaging assay, we used the fluorescent *nuIs25* (KP1148) strain in a *lite-1(ok530)* background. The *lite-1(lf)*, background was chosen to simplify the immobilization process by reducing the reactions of *C. elegans* animals to 488nm light. Once each day, we captured fluorescent images of glutamatergic neuronal cell bodies in the head of immobilized animals (**Fig. 5B**). We compared the fluorescent intensity of GLR-1::GFP in the soma of the AVA neurons in the worm each day by normalizing to the overall average on Day 1 (**Fig. 5C**). SAW immobilization facilitated the tracking of fluorescent intensity of the same individual worms over three days. As a comparison, we imaged control animals immobilized with sodium azide. Animals did not recover well after sodium azide exposure in the device, so we did not track single animals over multiple days. Instead, we imaged representative animals corresponding to the same age as their SAW immobilized counterparts (**Fig. 5B**). Comparing between SAW and control animals did not show a significant difference seen in the daily fluorescent intensity (**Fig. 5C**). Between 0 hours and 24 hours, SAW animals showed an average decrease of 38%, while representative control animals showed an average decrease of 41%. Between 24 hours and 48 hours, fluorescent intensity in SAW animals decreased a further 18%. Interestingly, the representative controls showed a 19% increase in average intensity. This difference is most likely due to the wide range of intensity between different individual animals which was seen both in SAW and control tests.

## Conclusion

This microfluidic SAW device provides a novel non-contact method for repeated, reversible, and temporally limited fast immobilization of individual *C. elegans* animals. By applying SAW in a precise and controlled manner to a simple, PDMS channel, we can use an acoustic approach to immobilize animals for a short period of time, allowing for high resolution fluorescent imaging. This enables the longitudinal analysis by fluorescence microscopy of single neurons of individual animals and their swimming behavior as they age in the same device. Here, we illustrate a proof of concept by imaging the glutamate receptor GLR-1 tagged with GFP in individual animals and tracking the fluorescent intensity change and bending frequency of animals over a 48-hour period. Our approach can accommodate longitudinal analysis of worms with improved precision by tracking a single animal over multiple days. In comparison, performing similar analysis by using different sets of animals may result in inconsistent measurements (**Fig. 5C**).

Other microfluidic methods to immobilize *C. elegans* have previously been reported, such as tapered channels^20^, flexible membranes^21–23^ CO_2_ introduction^23^, cooling^24^ and heating^25^. However, they suffer from drawbacks such as geometry constraints and significant environmental changes that have the potential to induce stress signaling in animals that could interfere with analysis. Our device facilitates an acoustic immobilization method with a standard channel geometry and a minimal environmental change, allowing for rapid switching between an immobilized and free-swimming state. The simple design of the chip lends itself to straightforward fabrication using standard photolithography techniques. Moreover, devices are indefinitely reusable (>50 cycles tested).

There are certain limitations to consider when using this immobilization method. Although SAW can immobilize worms quickly (~1 min), maintaining the immobilized state for multiple minutes tended to prevent full recoveries. Therefore, if longer-term immobilization is required, other methods may be more suitable. However, for rapid high-resolution imaging and dynamic behavior screening, our device is an ideal candidate.

In the future, the device can be further improved by characterizing its compatibility with long working distance objectives and more powerful microscopes to achieve better clarity, particularly at synaptic resolutions. Additionally, an automatic worm loading system to ensure a single worm is quickly and precisely inserted into the immobilization channel can be integrated into the PDMS chamber. The conversion from high to low SAW for microscopy immobilization could be automated based on live tracking of swimming. Environmental control channels could also be added to the system, allowing for precise inflows for food and chemical delivery. Overall, by allowing a combined analysis of microscopy-based cellular signaling and behavior, our device shows promise for use in future applications for multiplexed *C. elegans* aging studies.

## Acknowledgements

The microfluidic devices were fabricated in the JILA clean room at University of Colorado Boulder. Wild-type *C. elegans* animals and worm culturing media were provided by the Ding Xue Lab at University of Colorado Boulder. Some strains were provided by the CGC, which is funded by NIH Office of Research Infrastructure Programs (P40 OD010440). This work was supported by startup fund and W.M. Keck medical research grant to X.Ding, a CCTSI New Methods Development pilot grant shared by F. Hoerndli and X. Ding and part of the CCTSI NIH-NCATS CTSA Grant (UL1-TR002535-01), and internal funding from Colorado State University College of Veterinary Medicine and Biomedical Sciences to F. Hoerndli.

## Supplementary Figures

**Supplementary Figure 1:**
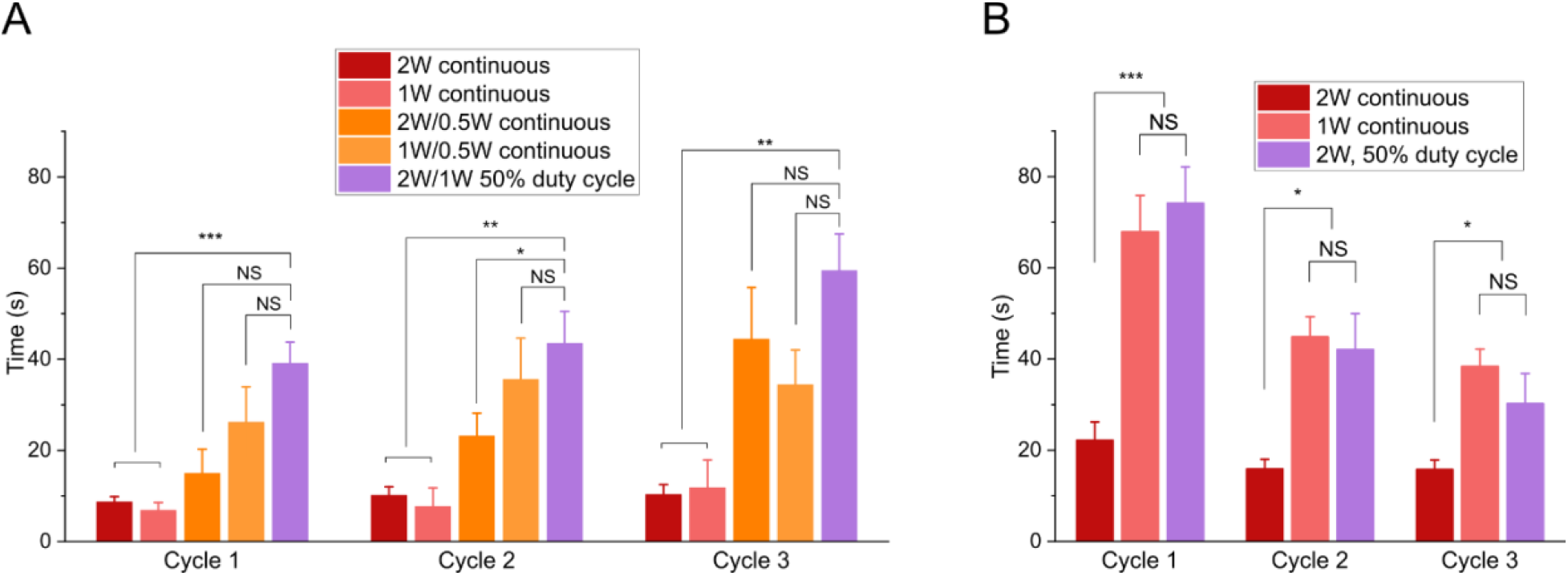
Characterization of worm immobilization and recovery using SAW device. **A,** Time required to immobilize animals using different SAW power levels and duty cycles. **B,** Total immobilization time after SAW without (red) or with (orange, magenta) a low power hold step. All conditions led to animal recovery after SAW and immobilization, Data are mean ± s.e.m. For each power level, at least 9 animals were tested for 3 cycles each. *P< 0.05, **P<0.01, ***P< 0.001, two-sided, unpaired t-test.

**Supplementary Figure 2:**
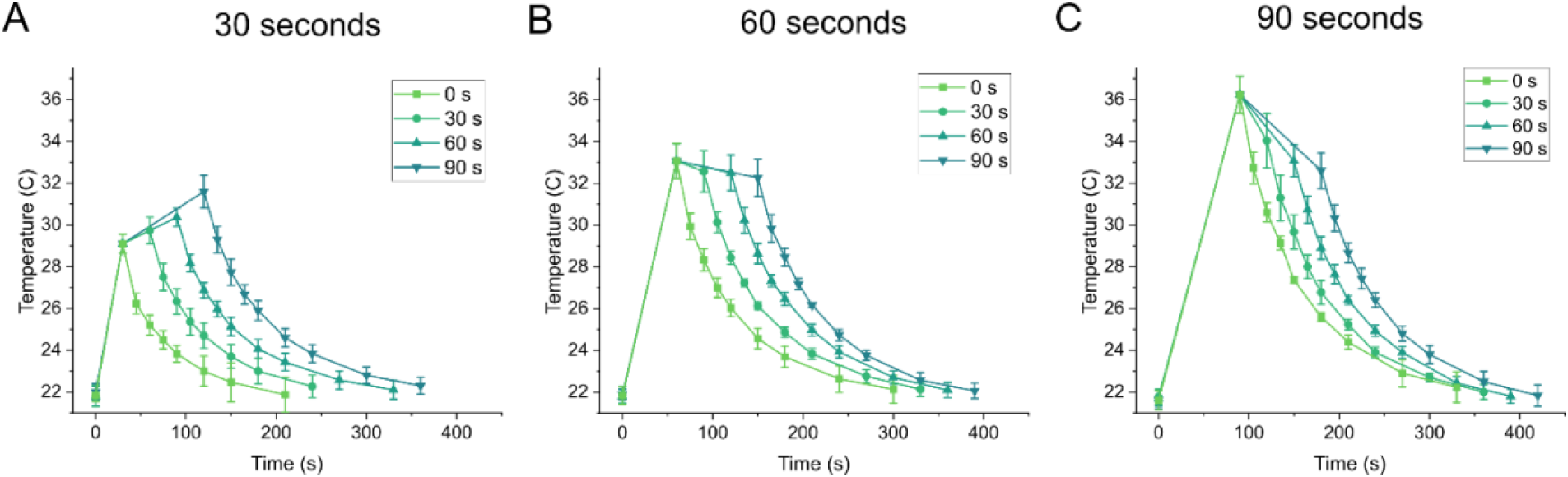
Temperature characterization of device showing initial high power SAW (2 W-50% duty cycle for **a,** 30 seconds, **b,** 60 seconds, and **c,** 90 seconds and corresponding low power SAW (1 W-50% duty cycle) hold times. Device cooling after SAW was turned off was also tracked. Data are mean ± s.e.m. Three individual experiments were performed for each temperature curve.

**Supplementary Figure 3:**
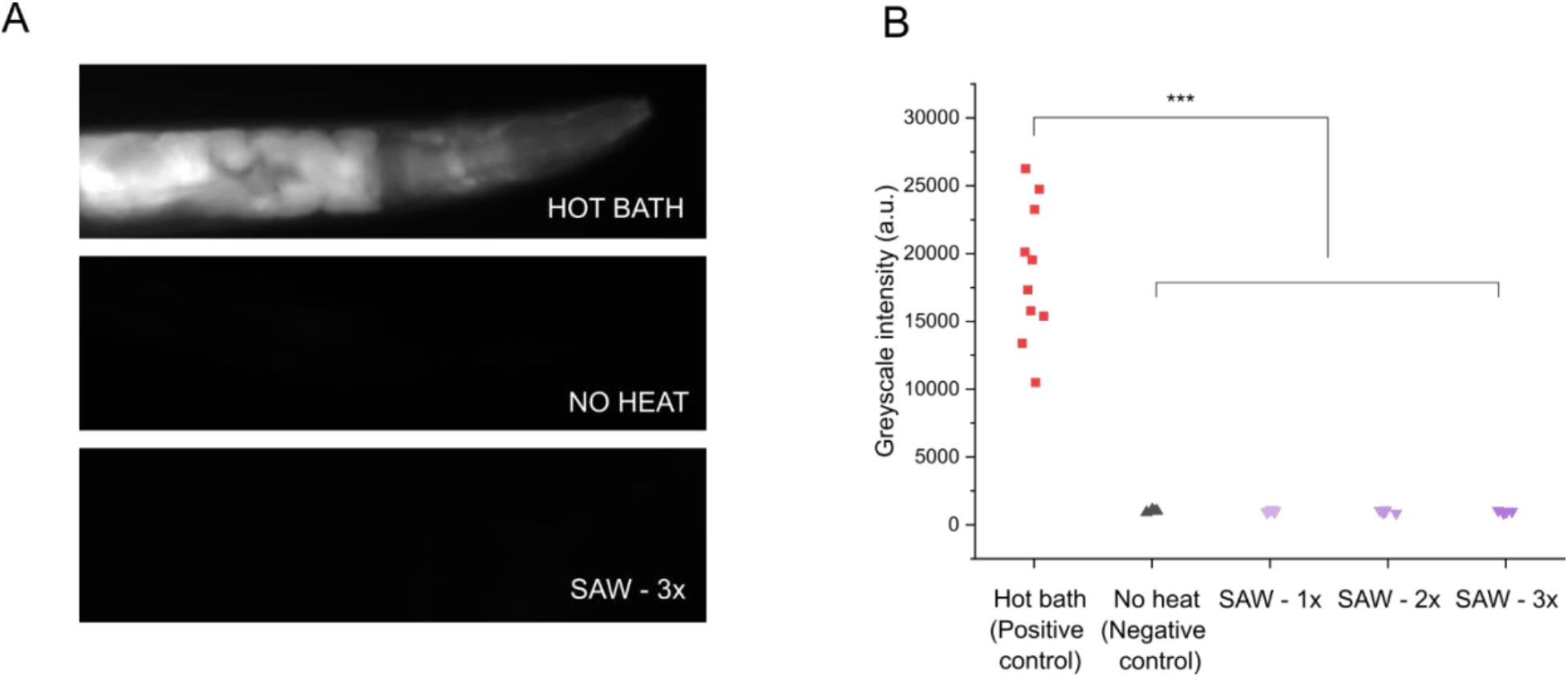
Heat shock characterization. **A,** Images of representative *ST66* animals 1 hour after (i) positive control – 1 hour hot bath at 32° C, (ii) negative control – no heat applied, and (iii) 3x SAW immobilization cycles. **B,** Greyscale intensity after measurements comparing positive and negative controls with 1, 2, and 3 SAW immobilization cycles. Data are individual animal measurements. At least 10 individual animals were imaged for positive control, and 5 individual animals were imaged for all other experiments. ***P< 0.001, two-sided, unpaired t-test.

